# Hurricane-induced population decrease in a Critically Endangered long-lived reptile

**DOI:** 10.1101/2021.07.06.451308

**Authors:** Matthijs P. van den Burg, Hannah Madden, Timothy P. van Wagensveld, Erik Boman

**Affiliations:** IUCN SSC Iguana Specialist Group, Gland, Switzerland; BioCoRe S. Coop. Calle Villagarcía 6, 28010, Madrid, Spain; Caribbean Netherlands Science Institute, P.O. Box 65, St. Eustatius, Caribbean Netherlands; NIOZ Royal Netherlands Institute for Sea Research, and Utrecht University, P.O. Box 59, 1790 AB Den Burg, Texel, the Netherlands; Reptile Amphibian Fish Research the Netherlands, Nijmegen, The Netherlands; St. Eustatius National Park Foundation, St. Eustatius, Caribbean Netherlands

**Keywords:** Abundance, Caribbean, Detection probability, Disturbance event, Habitat, Hurricane, *Iguana delicatissima*, Monitoring, Site occupancy, St. Eustatius

## Abstract

Catastrophic events, like hurricanes, bring lethal conditions that can have population-altering effects. The threatened Caribbean dry forest occurs in a region known for its high-intensity hurricane seasons and high species endemism, highlighting the necessity to better understand hurricane impacts as fragmentation and clearing of natural habitat continues. However, such studies remain rare, and for reptiles are mostly restricted to *Anolis*. Here we used single-season occupancy modeling to infer the impact of the intense 2017 Atlantic hurricane season on the critically endangered Lesser Antillean Iguana, *Iguana delicatissima*. We surveyed 30 transects across eight habitats on St. Eustatius during 2017-2019, which resulted in 344 individual surveys and 98 iguana observations. Analyses of abundance and site occupancy indicated both measures for 2018 and 2019 were strongly reduced compared to the pre-hurricane 2017 state. Iguanas at higher elevations were affected more profoundly, likely due to higher wind speeds, tree damage and extensive defoliation. Overall, our results indicate a decrease in population estimates (23.3-26.5%) and abundance (22-23.8%) for 2018 and 2019, and a 75% reduction in the number of opportunistic sightings of tagged iguanas between 2017-2018. As only small and isolated *I*. *delicatissima* populations remain, our study further demonstrates their vulnerability to stochastic events. Considering the frequency and intensity of hurricanes are projected to increase, our results stress the urgent need for population-increasing conservation actions in order to secure the long-term survival of *I*. *delicatissima* throughout its range.

## Introduction

Hurricanes bring lethal conditions to populations in their path. These events can either directly affect natural populations, through e.g., mortality (Reagan 1991; Wiley and Wunderle 1993; Spiller et al. 1998; Behie and Pavelka 2005; Morcilo et al. 2020; Marroquin-Paramo et al. 2021), or indirectly e.g., when food presence and abundance is impacted (Cely 1991). Selective mortality sweeps have been shown to drive both evolution (Donihue et al. 2018) and community diversity (Johnson and Winker 2010; Meléndez-Vazquez et al. 2019). Additionally, catastrophic winds can aid dispersal of species (e.g., Censky et al. 1998). Although studies on the effects of hurricanes are rising given their projected increase (Webster et al. 2005), such studies are dependent on pre-hurricane data and hurricane trajectory, and are thus lacking for many species.

From a conservation perspective, the population-diminishing threat of hurricanes is especially of concern for closed island populations, which often represent restricted-range and endemic species. As natural habitat disappears and fragmentation continues to worsen (Jacobson et al. 2019; Powers and Jetz 2019), understanding how stochastic events affect remaining populations is crucial. In a rare long-term study on an endangered Bahamian woodpecker (*Melanerpes superciliaris nyeanus*), Akresh et al. (2020) demonstrate that the population experienced significant declines as a result of various hurricanes but was able to recover after several years, however increased future hurricane frequency could prevent this. Contrarily, a study on a small, 350-individual, population of *Cyclura nubila* revealed that the passing of Hurricane Ivan (2004) had no effect on its size (Beovides-Casas and Mancina 2006), demonstrating the resilience of large rock iguanas to catastrophic events. Overall, studies concerning the effect of hurricanes on reptile species have mainly focused on small-size species (e.g., *Anolis*: Spiller et al. 1998; Schoener et al. 2001; Losos et al. 2003; Donihue et al. 2018, 2020; Dufour et al. 2019; Rabe et al. 2020). Insular iguanas are highly endangered ecological keystone species, and generally are the largest native remaining terrestrial species (IUCN 2020). As Caribbean hurricane frequency is projected to increase (Bender et al. 2010), understanding the impacts of hurricanes on iguana populations is important and is frequently highlighted but rarely quantified (Alberts 2004; Fogarty et al. 2004; Powell 2004; Pasachnik et al. 2012; but see Hayes et al. 2004, 2016; Beovides-Casas and Mancina 2006).

Among Caribbean Iguanids, the conservation status of the Lesser Antillean iguana (*Iguana delicatissima*) has experienced the most rapid deterioration (van den Burg et al. 2018a). Its pre-Colombian range has to date decreased by >80%, predominantly due to hybridization with *Iguana iguana* lineages (Vuillaume et al. 2015; Pounder et al. 2020), a process that was recently initiated on the last remaining ≥2 km^2^ islands where *I*. *delicatissima* occurs (van den Burg et al. 2018b, 2020; B. Angin pers. comm.). Left uncontrolled, the genetic loss of these populations will lead to a >99% decrease in their native range (van den Burg et al. 2018a). Critically, the five remaining populations that are free of nonnative iguanas occur on <2 km^2^ islets with low estimated population sizes (only Petite Terre >1,000 individuals; Angin 2017; Pounder et al. 2020). Beyond the spread of *I*. *iguana* (Knapp et al. 2020), other threats come from habitat destruction, pets and free-roaming livestock, as well as other anthropogenic causes (e.g., vehicle-caused mortality) (Debrot et al. 2013; Knapp et al. 2014, 2016; van den Burg et al. 2018c). Although hurricanes are cited as a potential threat to population survival (e.g., Fogarty et al. 2004), only two studies have addressed this topic. Namely, Knapp and Valeria (2008) provide some insight through an observational account of nest site degeneration and iguana mortality caused by Hurricane Dean (2007) on Dominica. Focusing on Petite Terre (Guadeloupe), a second study provides insight into a likely decrease in population size due to Hurricane Hugo (1995; Lorvelec et al. 2004). However, since the monitoring methodology has been questioned (Breuil and Ibéné, 2008; B Angin pers. comm.), no conclusions can be drawn. This study also predated new analytical tools and updates (e.g., Fiske and Chandler 2011).

Within the northern Lesser Antilles, St. Eustatius supports one of the last remaining *I*. *delicatissima* populations. This population suffered a major decline during the 1990s, in part due to consumptional demand on neighboring St. Maarten (Debrot et al. 2013), and is currently threatened by ongoing hybridization with *I*. *iguana* (van den Burg et al. 2018b). Despite the occasional arrival of stowaway iguanas, continuous monitoring has only resulted in 15 (Debrot et al. in review.) identified nonnative (pure or hybrid) animals, thus hybridization is believed to be minimal (unpublished data). The most recent estimates put the population size at below 500 individuals (Debrot et al. 2013), which has presumably since decreased due to ongoing anthropogenic threats (van den Burg et al. 2018c). Additionally to threats common for the species (see above), the population on St. Eustatius faces death by entrapment in abandoned cisterns, invasive rats, and the spread of the nonnative invasive Mexican creeper or Corallita (*Antigonon leptopus*) (Fogarty et al. 2004; Debrot et al. 2013; van den Burg et al. 2018c). Although St. Eustatius fell within the trajectory of major 2017-season hurricanes (Fig. 1), an impact assessment for *I*. *delicatissima* population is lacking.

**Fig. 1.**
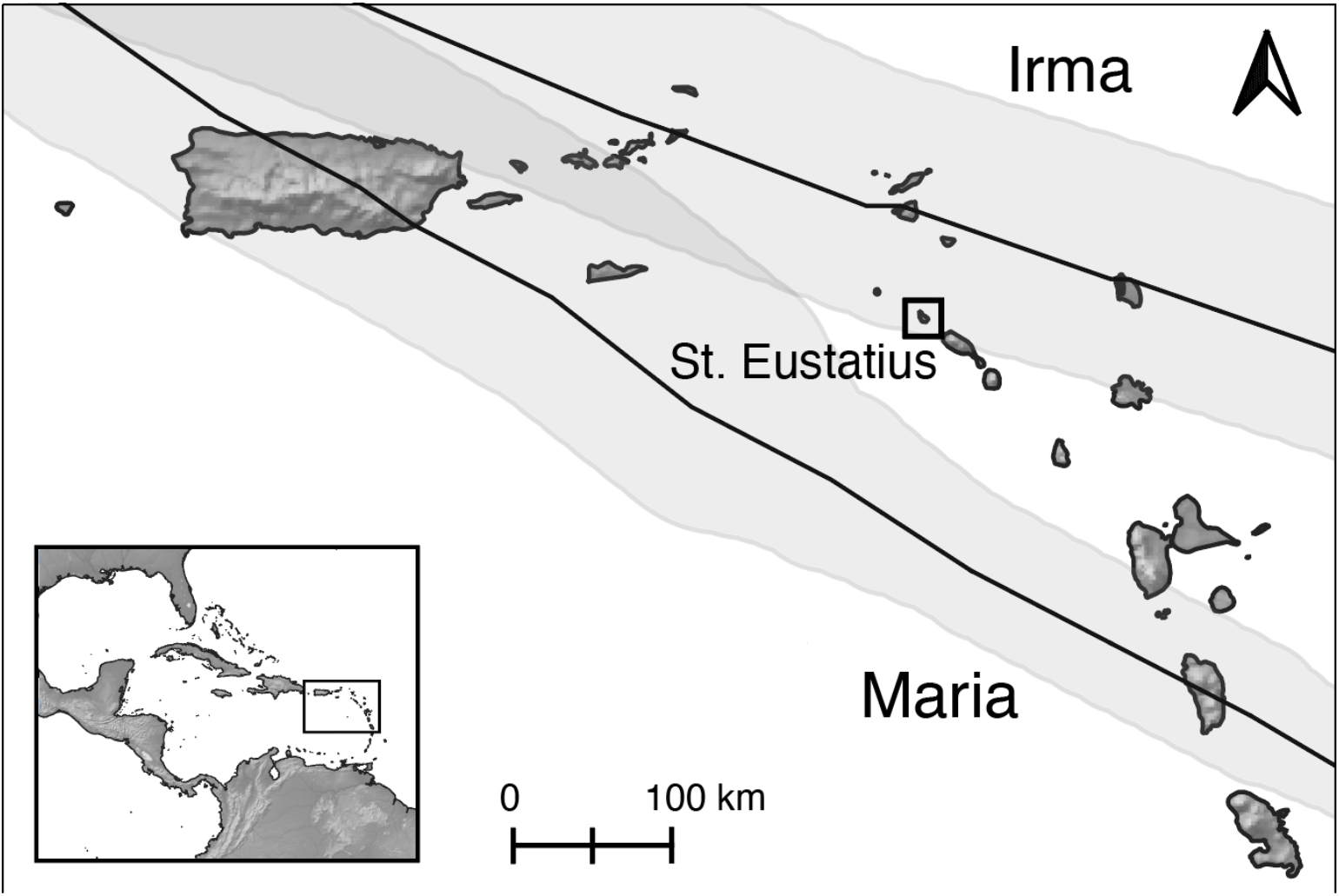
Best trajectory paths and regions with hurricane force winds (≥ 64kt) for Hurricane Maria and Irma in the northern Greater Caribbean region. Data from the National Hurricane Center.

The 2017 Atlantic hurricane season was categorized as hyperactive, with an accumulated cyclone energy score within the top ten since registration started (Schultz et al. 2018). Of the six 2017 hurricane-force systems, two closely passed St. Eustatius. On 6 September, the center of Hurricane Irma passed at ~54 km, and on 19 September, Hurricane Maria passed at roughly 110 km distance (Fig. 1; NOAA 2020). The goal of our study was to utilize iguana-survey data collected during 2017–2019 in order to assess the impact of the 2017 hurricane season on the St. Eustatius *I*. *delicatissima* population. To that end, we used island-wide transect survey data to estimate abundance, detection probability, and site occupancy per habitat type and year. Changes in occupancy, defined as the probability of a site being occupied (MacKenzie et al. 2002), provide a robust proxy for population declines, particularly for sparsely populated and cryptic species (Dénes et al. 2015; Beaudrot et al. 2016). We hypothesized that if population estimates were stable over this 3-year period, this would suggest that iguanas are resilient to catastrophic weather events. However, if population estimates varied, this would suggest that external factors (directly and/or indirectly) impacted iguana survival.

## Materials & Methods

### Study area

This study took place on St. Eustatius (17.49°N, −62.98°W), a small (21 km^2^) Leeward island within the northern Lesser Antilles (Fig. 1), with a human population of approximately 3,100 (Statistics Netherlands 2020). Habitat falls within the Caribbean dry forest and local climate is defined as Tropical Humid (Trewartha and Horn 1980), with over 90% of the island having a tropical savanna climate (<400 m) (De Freitas et al. 2012). Annual temperature fluctuates around 25–33 °C, with average annual rainfall around 1000 mm while the highest elevations receive <2,000 mm (Rojer 1997; van Andel et al. 2016). Geographically, the island consists of two mountainous areas that originate from volcanic activity and uplift on the northern and southern end. These areas are separated by a lower elevational plain where most anthropogenic activity occurs. Historic island-wide agricultural practices have downgraded the habitat quality which has partially recovered due to a significant decrease in such practices and conservation efforts, though habitats are affected still by grazers, erosion and invasive species (van Andel et al. 2016).

### Transects

Between 2017–2019, we conducted repeated count transect surveys to estimate site occupancy, detection probability and abundance of the *I*. *delicatissima* population (Royle and Nichols 2003). For this, we placed a total of 30 transects over the island, each 100 m in length, and visually surveyed up to 25 m on both sides (Fig. 2). Surveys were performed by minimally one observer with iguana-fieldwork experience, during hours of general iguana activity (Pasachnik et al. 2002), and were limited to non-rainy days given ectotherm activity. We recorded the date, survey duration (min.), elevation (m), number of observers, and observed number of iguanas. Due to limited capacity to conduct surveys, transects were surveyed a maximum of six times per year. Transects included 47% of present habitat types representing 78.3% of the island’s surface. Several habitat types were not surveyed due to access restrictions (e.g., *Bothriochloa* - *Antigonon*) or physical safety (e.g., *Antirhea* - *Coccoloba* mountains). Furthermore, the northern area was surveyed less often than other habitats due to logistics and access difficulty of the area, and given an oil transhipment company occupies 10–15% of the island’s surface area. Following De Freitas et al. (2012) nomenclature, surveys were performed in eight habitat types (see Fig. 2, Table 1): *Pisonia* - *Antirhea* mountains; miscellaneous urban/agricultural or disturbed land; *Pisonia* - *Justicia* hills; *Capparis* - *Pisonia* mountains; *Rauvolfia* - *Antigonon* mountains; *Coccoloba* - *Bothriochloa* cliffs; *Pisonia* - *Eugenia* mountains; *Chionanthus* - *Nectandra* mountains.

**Fig. 2.**
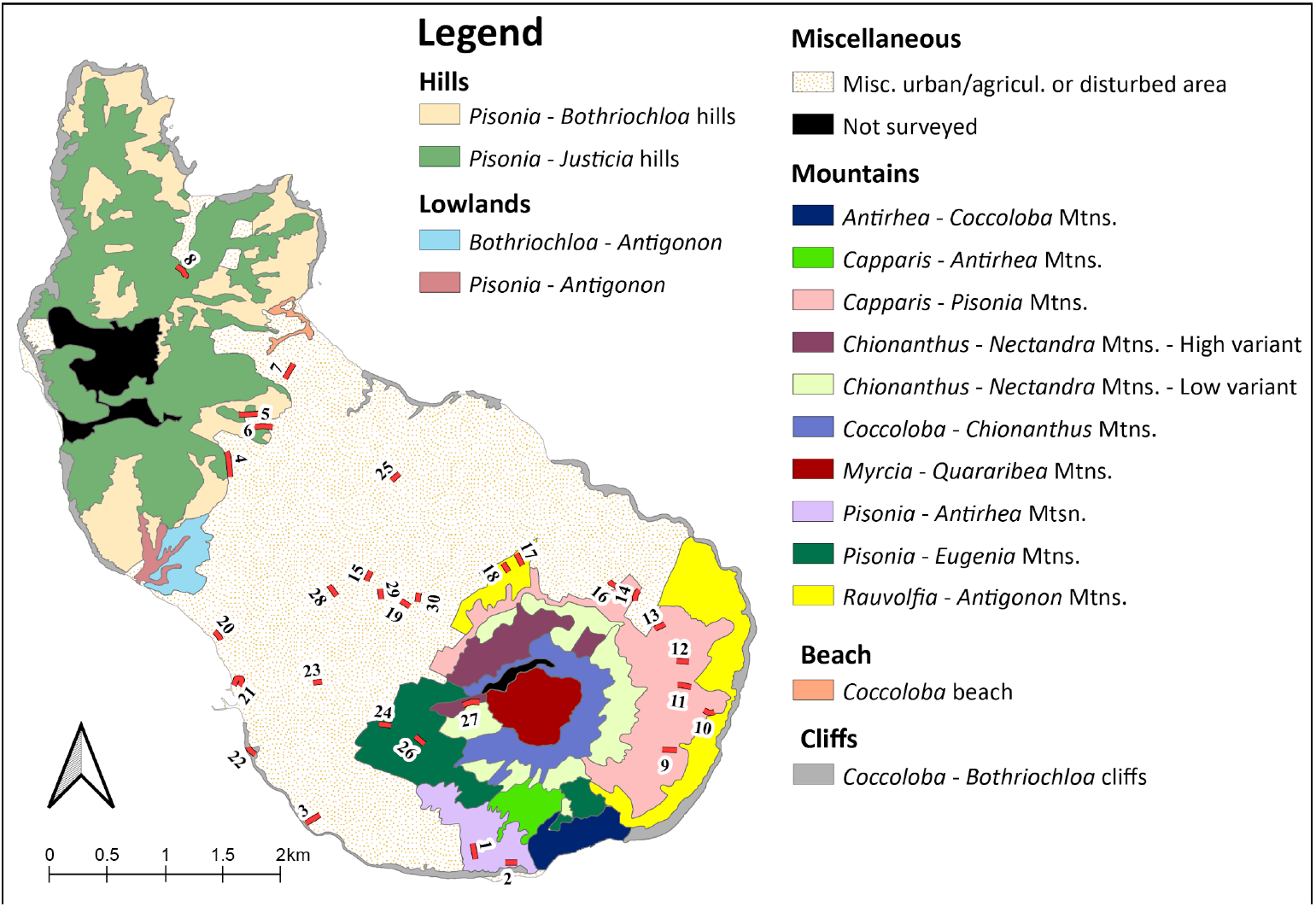
Map of St. Eustatius showing transect locations and habitat distribution. Habitat names following De Freitas et al. (2012).

**Table 1.** Summary of transect surveys conducted on St. Eustatius between 2017–2019. Habitat name following De Freitas et al. (2012). All transects constituted 100 m in length with a surveyed area of 500 m^2^.

| Habitat name | Transect nr. | Lat. | Lon. | Start elev. (m) |
| --- | --- | --- | --- | --- |
| *Pisonia* - *Antirhea* mountains | (1) | 17.4672 | -62.9681 | 69 |
|  | (2) | 17.4659 | -62.9643 | 41 |
| Misc. urban/agricultural or disturbed land | (3) | 17.4691 | -62.9808 | 12 |
|  | (4) | 17.4947 | -62.9867 | 59 |
|  | (5) | 17.4985 | -62.9835 | 53 |
|  | (7) | 17.5024 | -62.9825 | 25 |
|  | (15) | 17.4869 | -62.9759 | 93 |
|  | (16) | 17.4868 | -62.9573 | 117 |
|  | (19) | 17.4847 | -62.9728 | 141 |
|  | (20) | 17.4831 | -62.9879 | 14 |
|  | (21) | 17.4797 | -62.9860 | 25 |
|  | (23) | 17.4794 | -62.9801 | 87 |
|  | (25) | 17.4945 | -62.9742 | 48 |
|  | (28) | 17.4859 | -62.9783 | 91 |
|  | (29) | 17.4856 | -62.9749 | 114 |
|  | (30) | 17.4862 | -62.9721 | 123 |
| *Pisonia* - *Justicia* hills | (6) | 17.4994 | -62.9844 | 66 |
|  | (8) | 17.5098 | -62.9900 | 70 |
| *Capparis* - *Pisonia* mountains | (9) | 17.4742 | -62.9518 | 79 |
|  | (10) | 17.4774 | -62.9499 | 50 |
|  | (11) | 17.4790 | -62.9508 | 72 |
|  | (12) | 17.4806 | -62.9510 | 80 |
|  | (13) | 17.4837 | -62.9527 | 104 |
|  | (14) | 17.4862 | -62.9547 | 99 |
| *Rauvolfia* - *Antigonon* mountains | (17) | 17.4892 | -62.9642 | 104 |
|  | (18) | 17.4884 | -62.9654 | 117 |
| *Coccoloba* - *Bothriochloa* cliffs | (22) | 17.4745 | -62.9852 | 12 |
| *Pisonia* - *Eugenia* mountains | (24) | 17.4764 | -62.9748 | 149 |
|  | (26) | 17.4754 | -62.9726 | 190 |
| *Chionanthus -*<br>*Nectandra* mountains | (27) | 17.4780 | -62.9670 | 386 |

### Opportunistic surveys/sightings

Since 2015, subadult and adult iguanas have been tagged with colored glass beads to facilitate subsequent noninvasive tracking and observations of individuals using unique color combinations (Binns and Burton 2007). To date, 540 unique bead codes have been assigned, and an additional 96 iguanas have been tagged only with PIT tags. This methodology has allowed for opportunistic data collection of the geographic locations of bead-tagged iguanas. We utilized these data as an additional indicator for variation in population size.

### Data analyses

As our dataset contained a large number of non-detections and small counts of iguanas, we chose an *N*-mixture model approach over distance sampling (Royle 2004). Models were based on four assumptions: (i) the iguana population in each site was closed to colonization and extinction during the sampling period; (ii) iguanas were either detected or not detected during surveys; (iii) iguanas were not falsely detected during surveys; and (iv) iguana detections at each site were independent (Murray and Sandercock 2020). We followed the methods described by Madden et al. (2021). Briefly, we used likelihood-based single-season occupancy models (MacKenzie et al. 2017), using the R packages “wiqid” and “unmarked” (MacKenzie et al. 2002; Fiske and Chandler 2011), to estimate site occupancy (ψ) in relation to habitat and elevation, while accounting for detectability (*p*). We modeled detection probability and occupancy probability to estimate iguana abundance (λ), and also tested the influence of independent variables on abundance estimates, including quadratic terms. Resulting models were ranked using Akaike’s Information Criterion, where the detection model with the lowest AICc is best-fitting [AICc; Burnham and Anderson (2002)], while correcting for small sample size. Finally, we tested for overdispersion by assessing the goodness-of-fit of the most parsimonious model in each year (MacKenzie and Bailey 2004) with the “mb.gof.test” function in the package “AICcmodavg” using 1,000 simulations, which calculates a Pearson’s chi-square fit statistic from the observed and expected frequencies of detection histories for a given model. All analyses were performed in the R environment v3.5.1 (R Core Team 2019).

## Results

### Pre-hurricane survey occupancy, detection and abundance estimates

We conducted 180 surveys between February and April 2017, prior to hurricanes Irma and Maria, during which we detected 58 individual iguanas. Average transect time was 32 ±14 minutes. As results for both R packages provided identical patterns we only report “wiqid” results here, and “unmarked” results are presented in the Electronic Supplementary Information.

Occupancy probability was 100% (range 0–100%), and iguana occupancy and abundance increased with elevation (Figs. 3–4) as well as with survey effort. Occupancy estimates were generally consistent across the habitats sampled (Table 2). Mean detection probability (*p*) per transect was 0.19 (range 0.14–0.26), which increased to 0.27 when an iguana had been previously detected. The effects of predictor variables ‘survey time’ and ‘week’ on detection and occupancy estimates were negligible (**Δ**AIC*_c_* <2). We observed a lack of fit of the null model (without covariates; *X*^2^ = inf, p = 0, c-hat (ĉ) = inf). Estimated mean individual abundance (λ) was 5.91 (± SE 0.01). When habitat and elevation were included as response variables, λ estimates were 1.52 ± 0.65 (Fig. 5), which translates to a population estimate across the entire study area (13.5 ha) of 1,364 iguanas. Lowest iguana abundance was found in *Coccoloba* - *Bothriochloa* cliff habitat, and highest in *Chionanthus* - *Nectandra* mountain habitat.

**Fig. 3.**
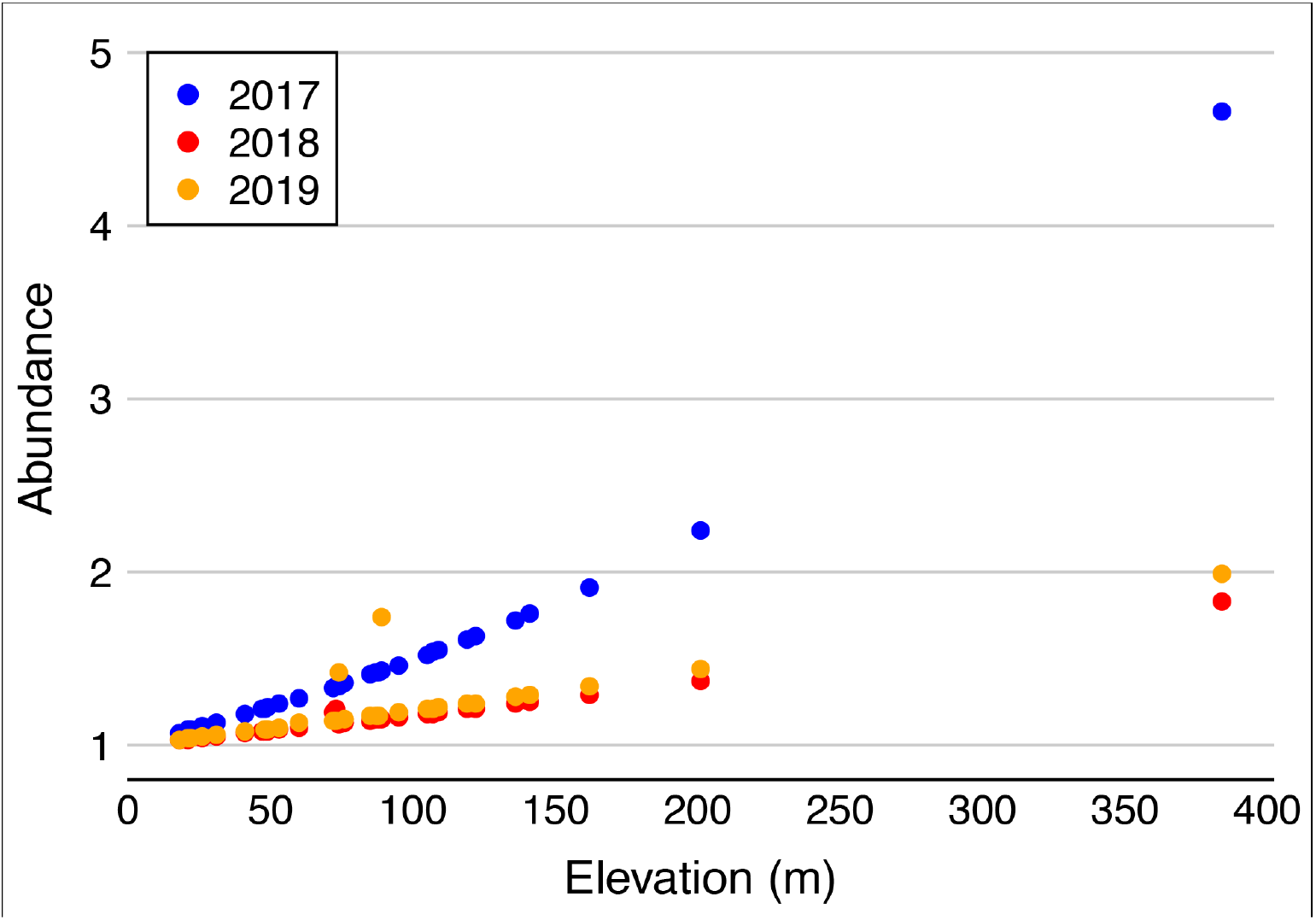
The effect of elevation on abundance (lambda) estimates of *Iguana delicatissima* based on transect surveys conducted on St. Eustatius in 2017 (pre-hurricane), 2018 and 2019 (post-hurricane).

**Fig. 4.**
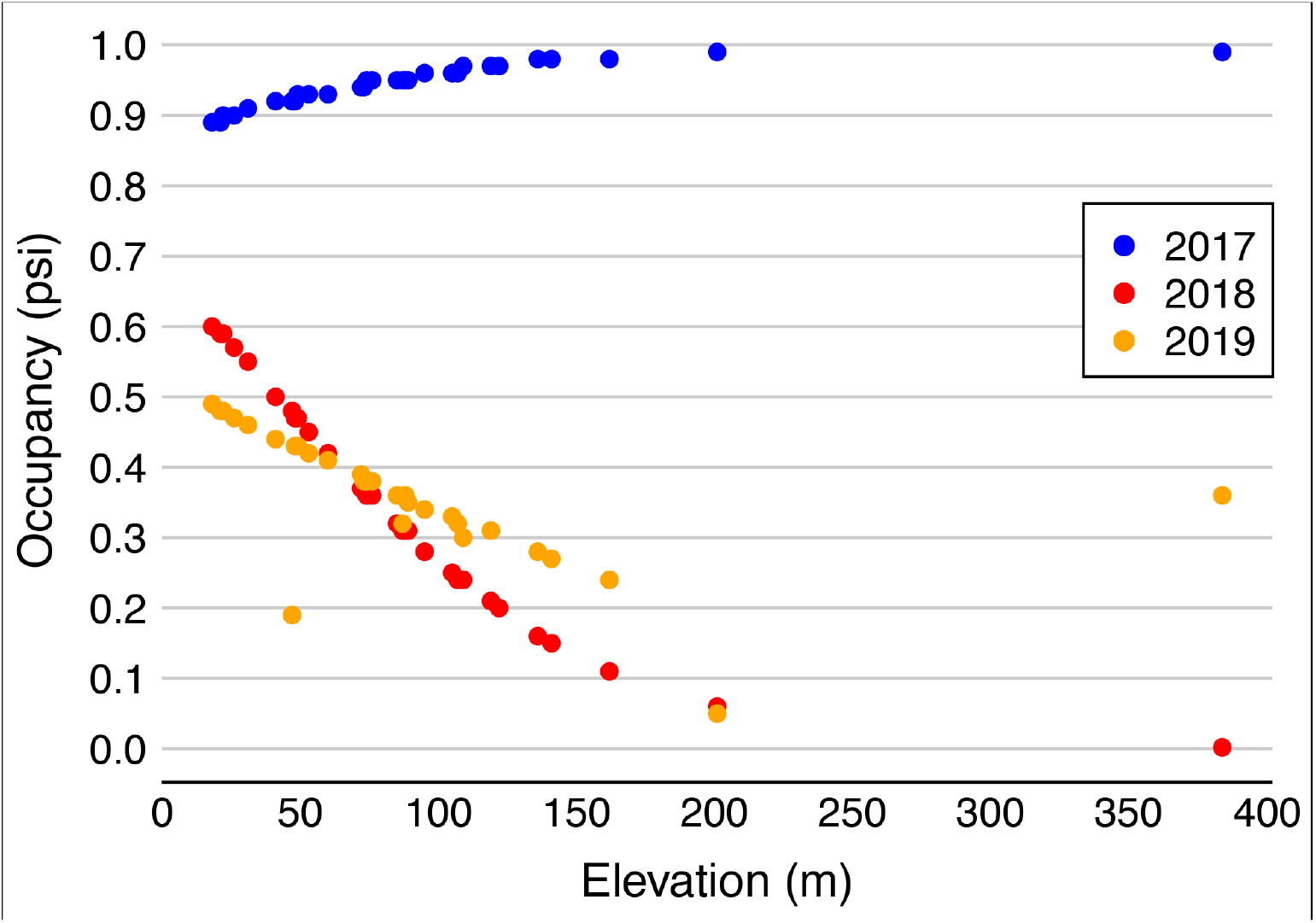
The effect of elevation on site occupancy estimates of *Iguana delicatissima* based on transect surveys conducted on St. Eustatius in 2017 (pre-hurricane), 2018 and 2019 (post-hurricane).

**Fig. 5.**
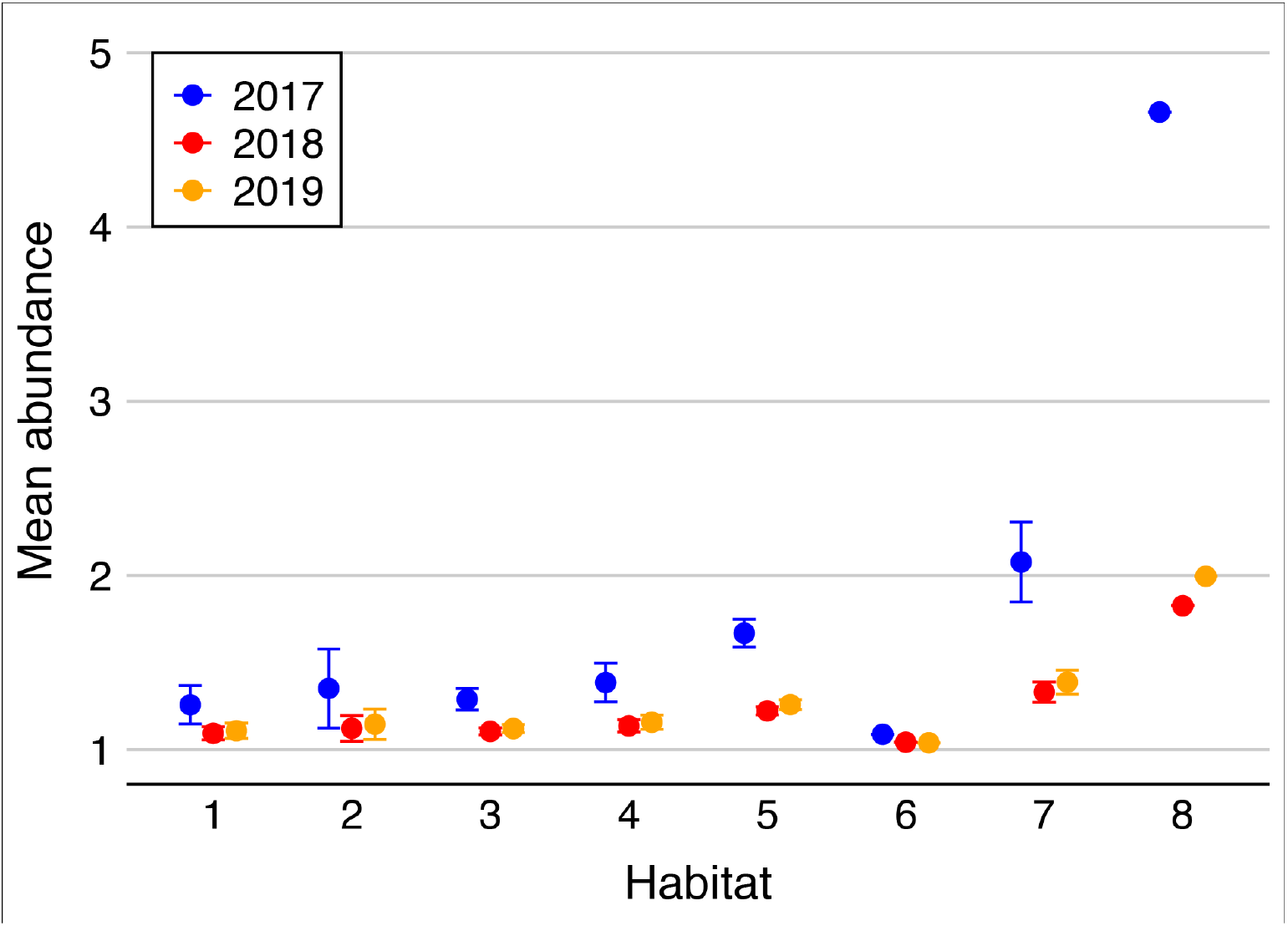
Mean iguana abundance estimates per habitat type for 2017–2019, with standard deviations bars. Habitat following De Freitas et al. (2012): 1 = *Pisonia* - *Antirhea* mountains; 2 = miscellaneous urban/agricultural or disturbed land; 3 = *Pisonia* - *Justicia* hills; 4 = *Capparis* - *Pisonia* mountains; 5 = *Rauvolfia* - *Antigonon* mountains; 6 = *Coccoloba* - *Bothriochloa* cliffs; 7 = *Pisonia* - *Eugenia* mountains; 8 = *Chionanthus* - *Nectandra* mountains.

**Table 2.** Overview of 2017–2019 site occupancy estimates per habitat of *Iguana delicatissima* on St. Eustatius. Data represents the probability of a site to be occupied (Ψ), as well as confidence intervals (CI). Habitat following De Freitas et al. (2012): 1 = *Pisonia* - *Antirhea* mountains; 2 = miscellaneous urban/agricultural or disturbed land; 3 = *Pisonia* - *Justicia* hills; 4 = *Capparis* - *Pisonia* mountains; 5 = *Rauvolfia* - *Antigonon* mountains; 6 = *Coccoloba* - *Bothriochloa* cliffs; 7 = *Pisonia* - *Eugenia* mountains; 8 = *Chionanthus* - *Nectandra* mountains.

| Habitat | 2017 |  | 2018 |  | 2019 |  |
| --- | --- | --- | --- | --- | --- | --- |
| | $\psi$ | CI | $\psi$ | CI | $\psi$ | CI |
| 1 | 0.98 | <0.001–1.00 | 0.37 | 0.07–0.81 | 0.37 | 0.12–0.71 |
| 2 | 0.98 | <0.001–1.00 | 0.35 | 0.10–0.73 | 0.37 | 0.16–0.63 |
| 3 | 0.99 | <0.001–1.00 | 0.34 | 0.11–0.66 | 0.37 | 0.19–0.59 |
| 4 | 0.99 | <0.001–1.00 | 0.32 | 0.11–0.64 | 0.36 | 0.18–0.61 |
| 5 | 0.99 | <0.001–1.00 | 0.31 | 0.09–0.66 | 0.36 | 0.13–0.67 |
| 6 | 0.99 | <0.001–1.00 | 0.29 | 0.06–0.72 | 0.36 | 0.09–0.76 |
| 7 | 0.99 | <0.001–1.00 | 0.28 | 0.04–0.78 | 0.36 | 0.06–0.83 |
| 8 | 1.00 | <0.001–1.00 | 0.26 | 0.02–0.84 | 0.35 | 0.04–0.88 |

### Post-hurricane survey occupancy, detection and abundance estimates

Post-2017, we surveyed identical transects between July–December 2018 and March–September 2019. Overall, 164 transects were performed during which we detected 13 iguanas in 2018 and 27 iguanas in 2019. Average transect time was 19 ± 8 minutes.

Occupancy probability was 32.5% (range 11.8–63.4%) in 2018, and 36.7% (19.2–58.5%) in 2019. Iguana occupancy decreased with increasing elevation in 2018 and 2019 (Fig. 3), but increased with survey effort; up to four repeats in 2018 and three in 2019. Occupancy estimates were lower than those from 2017 but generally consistent across habitats (Table 2). *p* was 0.41 (range 0.14–0.74) in 2018, and 0.67 (0.39–0.87) in 2019, decreasing to 0.28 (0.07–0.67; 2018) and 0.36 (0.14–0.66; 2019) when an iguana had been previously detected. λ was 5.89 ± 0.02 in both 2018 and 2019. Considering habitat and elevation, λ estimates were 1.16 ± 0.15 in 2018 and 1.19 ± 0.18 in 2019 (Fig. 5), translating to population estimates of 1,046 and 1,003 iguanas, respectively; declines of 23.3% and 26.5% compared to 2017. The covariate ‘survey time’ from the most parsimonious model had a positive effect on detection probability (0.80 ± 0.49) in 2018, however we observed a lack of fit of this model (‘survey time’; *X*^2^ = inf, p = 0, ĉ = inf). The effect of covariates on detection and occupancy estimates in 2019 were negligible (**Δ**AIC*_c_* <2). There was no evidence for lack of fit in 2019 (*X*^2^ = 33.86, p = 0.66, ĉ = 0.62). Habitat abundances were lower in 2018 and 2019 compared to 2017, with only abundance range overlap in the *Pisonia* - *Antirhea* mountain and miscellaneous urban/agricultural or disturbed land habitat types (Fig. 5); compared to 2017, average decline in iguana abundance among habitats was 23.8% and 22.0% for 2018 and 2019, respectively. Overall, a two-way additive ANOVA indicated no significant difference of iguana abundance among years (F_2,14_ = 3.49, *p =* 0.059), and a significant difference between habitats (F_7,14_ = 3.76, *p =* 0.017).

### Bead-tag data

Opportunistic sighting data were collected during 76, 78 and 56 days in 2017, 2018 and 2019, respectively. Assessment of the geographic distribution of those data indicates both time and geographic effort was comparably equal for 2017 and 2018, allowing a direct comparison (see Electronic Supplementary Information), though not for 2019. Of the 145 uniquely bead-tagged iguanas observed in 2017, 36 (24.8%) were observed during 2018.

## Discussion

Our study demonstrates that multiple *I*. *delicatissima* population parameters decreased following the high-intensity hurricane year of 2017, with a decrease in estimates of population size, abundance, site occupancy, and opportunistic sightings of tagged iguanas. We did not observe a clear boomerang effect, as 2018-2019 parameters were mostly stable. These data are important to consider from a conservation and long-term stability perspective, given the small sizes of remaining populations of this highly endangered species. Overall, we present a rare effect-analysis of hurricane impact on a wild population of a large, long-lived reptile.

*Iguana delicatissima* inhabits most vegetation types within its native range, e.g., mangroves, high evergreen forest, dry shrublands, and gardens (for an overview see Henderson and Powell [2009]). On St. Eustatius, earlier studies found that iguana abundance was highest on vegetated slopes/cliffs and on the northwest flank of the dormant volcano where large estates are built within the tall evergreen forest (Fogarty et al. 2004; Debrot et al. 2013). Similarly, we found that iguana abundance was high in habitats on the western slopes of the volcano, though transects were not placed in gardens (Fig. 5). Thus, besides gardens, also the periferic habitats of this residential area are important iguana habitats. Indeed, Knapp and Perez-Heydrich (2012) found that biometric variables did not differ between iguanas living in- and outside of gardens. Furthermore, highest iguana abundance was found in the *Chionanthus* - *Nectandra* mountain habitat (De Freitas et al. 2012), at high (386m) elevation on the volcano, however this habitat was covered by a single transect, thus we lack a perspective of variation. Additionally, we found higher iguana abundances in the *Pisonia-Eugenia* mountain habitat, located on the lower/mid (150–200m) south-western slopes of the Quill. This habitat is described as a secondary forest and, according to van Andel et al. (2016), “has a much lower diversity, density and canopy height than the forest on the wetter slope”. Nevertheless, it is likely suitable and heterogenous habitat for iguanas and supports a variety of tree species including *Bourreria baccata, Pisonia subcordata, Guettarda scabra* and *Bursera simaruba* (van Andel et al. 2016) of which several are a known dietary source (Angin and Questel in prep.).

In contrast, *Rauvolfia - Antigonon* habitat is described as disturbed, regenerating shrubland with few trees, low floristic diversity and is covered extensively by an invasive vine (van Andel et al. 2016). Our results show that iguana abundance is surprisingly relatively high in this habitat type. We note that the patch covered by our survey efforts appears of higher quality than the remainder of this habitat type on the south-western side of the island, which is heavily grazed by free-roaming herbivores (van Andel et al. 2016; Madden 2020). Namely, the included patch has larger trees, is less degraded, and lies adjacent to secondary roads and semi-urban development. Consequently, we believe that our abundance estimates only apply to the covered patch of *Rauvolfia - Antigonon* habitat.

Detection probability increased from 2017 to 2019 but was overall low (<0.5) across all years. Nevertheless, detection probability estimates were similar to/higher than those from comparable studies (Bock et al. 2016; Rivera-Milán and Haakonsson 2020). However, when an iguana was previously observed, detection probability was equal across years. We did not test for observer effort, which may account for unequal detection between years and should ideally be consistent (Burton and Rivera-Milán 2014).; for our surveys, average number of observers was constant over the years (between 2–2.4), and surveys were always led by an experienced iguana-fieldwork technician. Vegetation damage (e.g., broken branches) may have resulted in higher detection probability post-hurricane compared to pre-hurricane, though generally iguanas are harder to detect in forested areas than other habitats. We were unable to locate specific published studies on iguanas to support this, but reptile species overall are cryptic and considered difficult to study (e.g., Durso et al. 2011; Sewell et al. 2012).

Our study provides evidence that the St. Eustatius *I. delicatissima* population declined during 2017, during which two major catastrophic events hit the island; Hurricanes Maria and Irma. As behavioural data of iguanas during such events are absent, pinpointing the exact cause of decline is more challenging; how did iguanas die, and was that during or after the events? Post-event mortality, in the absence of sustained physical injury, could result from a lower abundance of food. Indeed, the majority of trees were defoliated due to high wind speeds and likely salt spray (Eppinga and Pucko 2018). However, as not all trees were defoliated, and iguanas are known to survive extended periods of drought (van Buurt 2010), we anticipate that post-event mortality due to starvation had a minor contribution to the observed declines. Instead, we hypothesize that hurricane conditions, either led to deadly physical injuries or translocated iguanas into the Caribbean Sea (see Censky et al. 1998). Whether non-deadly on-island translocation is possible is unknown, though as both abundances declined for all habitat and opportunistic sightings declined drastically, this possibility appears unlikely. Although challenging to collect, future data on iguana movement during hurricane events could provide more insight.

We found evidence for overdispersion when calculating site occupancy measures for 2017 and 2018, which resulted in inflated standard errors (Table 2). However, given the lack of significance of covariates overall (only ‘survey time’ increased detection probability in 2018), we chose to retain AICc model weights than pursue a quasi-likelihood theory (MacKenzie and Bailey 2004). We recognize the limitations of our study with regard to the assumptions of occupancy modeling (Murray and Sandercock 2020), however we are confident that we met most if not all of the assumptions. We assumed that iguana detection was independent between sites, given the species’ limited home range (van Wagensveld et al. in prep.), and that the sites surveyed were discrete patches of habitat (except one survey track in the *Chionanthus - Nectandra* mountains habitat). Nevertheless, we recognize that there may be more variation in the observed site occupancy data than expected by the models. Considering the already wide confidence intervals, however, we are of the opinion that the models presented offer a realistic representation of the pre-/post-hurricane population data.

Despite the small size of St. Eustatius, our results suggest that the 2017 hurricanes affected iguanas disproportionately across the island. Namely, although abundance and occupancy decreased island wide, iguanas at higher elevations suffered stronger declines (Fig. 3–4). This finding is corroborated by data from Eppinga and Pucko (2018) who assessed the impact of both 2017 hurricanes on the forests of St. Eustatius and found that a higher percentage of trees was snapped at higher elevations. Eppinga and Pucko (2018) further show that over 90% of all trees were defoliated for >75% and over 75% of trees lost primary branches, suggesting that Irma and Maria heavily affected arboreal species. Indeed, hurricane wind speeds are stronger at higher elevations (Holthuijsen et al. 2012), whereby our findings could be explained by increased mortality through physical injury during hurricane events, as well as subsequently through longer food unavailability (Cely 1991).

Understanding the natural movements of study species is important to consider in population assessments (Neilson et al. 2018). Both *Iguana* species are known to remain static throughout most of their life cycle, with movement mainly restricted to the hatchling life stage until settling in breeding populations, and by adult females while traveling to nesting sites (Burghardt and Rand 1985; Bock and McCracken 1988; Knapp et al. 2016). For *I*. *delicatissima*, the reproductive cycle of populations on arid islands is clearly defined with nesting mainly occurring in June-August (van den Burg et al. 2018a). Reproductive data for the St. Eustatius population remains limited but appears to occur in June-July (pers. obs. by authors), which was considered while planning field surveys. As no large communal nesting sites occur on St. Eustatius (as opposed to Dominica, Knapp et al. 2016), and small isolated nesting sites have been located around the island (Debrot et al. 2013; pers. obs. by authors), we suspect gravid females to undertake little movement and nest within or close to their home range. This is further strengthened by the restricted size and highly fragmented status of the population (Debrot et al. 2013; van den Burg et al. 2018b), with interaction observations being rare (pers. obs. by authors).

Despite over half of Iguanidae diversity occurring in the Greater Caribbean region, hurricane impacts on their populations have been rarely addressed. Beovides-Casas and Mancina (2006) performed a total of 60 surveys (12 each during 4 visits) of the *Cyclura nubila* population on Cayo Sijú (Cuba) in 2004, with the final visit post-Hurricane Ivan. Although Ivan passed at ~160 km distance with sea level rising by 2.5 meter, their data indicated the population size remained stable. In another study, Hayes et al. (2004) report local extinction of a *Cyclura rileyi* population from a small Bahamian cay following the passing of Hurricane Floyd (1999). However, it is important to note that the authors indicated only three animals were observed by themselves there, with a personal comment from the previous decade about 18 animals. Thus, the subsequent reference of population extirpation following a hurricane event (e.g., Hayes et al. 2016) without mention of these numbers and overall poor knowledge of the population in question appears unjustified. Additionally, Hayes et al. (2004) also report no decrease in adult iguanas from a nearby cay, with a large population. In our study, we found that multiple population parameters decreased after an intense hurricane season; a decrease in I) population size of the studied area (23.3–26.5%) and II) abundance (22–23.8%), as well as III) a 75% reduction in the number of opportunistic sightings of tagged iguanas between 2017–2018. However, as studies remain scarce, equally are generalizations and comparisons. Compared to *I*. *delicatissima*, *Cyclura* sp. are larger and more robust, mainly non-arboreal iguanas. Although pre-hurricane behavior for *I*. *delicatissima* is unknown, presumably lighter iguanas that remain arboreal during catastrophic weather events are more vulnerable than heavier, terrestrial iguanas. However, inundation might force *Cyclura* sp. to ascend trees in order to avoid being swept away given their range mostly includes low-elevation islets.

To evaluate the impact of catastrophic events on closed populations, consideration of these events’ timing compared to a species’ reproductive cycle is essential (Schoener et al. 2004). For St. Eustatius, both 2017 hurricanes passed as strong early storms, preceding the main hatching period of the local *I*. *delicatissima* population while most eggs were incubating and therefore presumably safe in the absence of inundation. Later-seasonal storms are expected to have a more severe impact, as hatchlings are vulnerable to high wind speeds, thus potentially losing most individuals of an entire generation. Indeed, Hayes et al. (2004) compared hatching percentage between subsequent years and reported an extremely small hatchling cohort after the passing of Hurricane Floyd, which hit the Bahamas during the hatching period of *C*. *rileyi*. No information is available about the percentage of hatched clutches. On St. Eustatius only some nesting occurs on the island’s lower elevations with the majority of nest sites found at elevations far higher than sea-level inundation (Debrot et al. 2013). Although data on egg survival and hatching success after inundation is known for some reptilians (Hsu et al. 2021), such data is missing for Iguanids.

Effect assessments of the 2017 Atlantic hurricane season are demonstrating its broad impact on ecosystems and their trophic levels; e.g., beach, mangrove and forest ecosystems (Liu et al. 2018; Barreto-Orta et al. 2019; Walker et al. 2019; Hall et al. 2020), salt- and freshwater fish communities (Meléndez-Vazquez et al. 2019; Neal et al. 2020), insect species and communities (Cabrera-Asencio and Meléndez-Ackerman 2021), birds (Palmer et al. 2018; Lloyd et al. 2019), and seas slugs (Middlebrooks et al. 2020). For St. Eustatius specifically, our data adds to the understanding of an ecosystem-wide hurricane impact during 2017 which, besides iguanas, affected both forests (Eppinga and Pucko 2018) and the populations of Bridled Quail-dove (Rivera-Milán et al. 2021) and Red-bellied racer (Madden et al. 2021). Importantly, these insights and novel risk assessments are demonstrating the high extinction risk that endemic, island species will face under projected climate change (Manes et al. 2021).

### Conservation

From a conservation perspective, the status of *I. delicatissima* is at its most benign, with only five populations not directly threatened by on-island hybridization, of which just a single island has a >200 population size; Petite Terre (Guadeloupe). Our study further highlights this vulnerability, as local extinction or population declines to unsustainable sizes could be the result of major hurricanes. Especially the extremely small population of Anguilla is vulnerable (Pounder et al. 2020), which currently occurs on a small low-level islet. There, the potential of temporarily holding iguanas to increase hurricane survival should be assessed, given adequate facilities can be utilized. On St. Eustatius, we found no post-hurricane population recovery (or boomerang effect), which could be linked to the small size of the existing population. Therefore, the projected increase in frequency and intensity of catastrophic hurricanes (Bender et al. 2010) is troubling as populations might be unable to recover between subsequent catastrophic events (McCoid 1996; Schriever et al. 2009; Selman 2015). However, we note that a short-term boomerang effect was likely absent given the apparent low fecundity on St. Eustatius, possibly due to the population’s low genetic diversity (van den Burg et al. 2018b), although a detailed study on fecundity still is absent. Overall, our study strengthens the need for population-increasing measures to be taken.

## Acknowledgments

We thank Adam Mitchell, Sarinda Westerhout, Joey Markx, Rupert Redan, and all STENAPA staff and volunteers that helped with data collection. Funding was provided through the Mohamed bin Zayed Species Conservation fund (172517158), International Iguana Foundation, and, the BEST 2.0 program.

## Electronic Supplementary Material

### List of figures and table in this document

**Figure S1.**
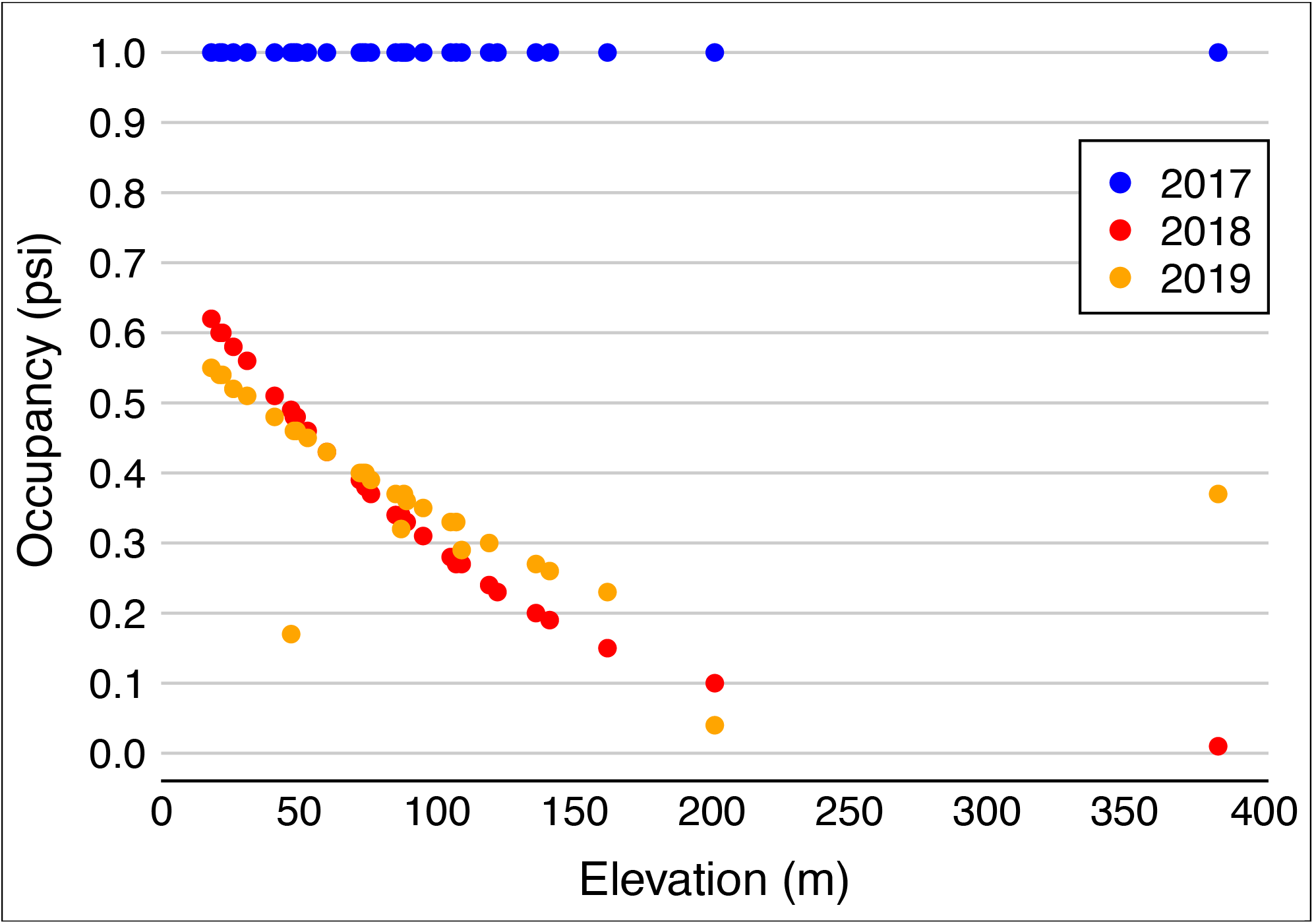
The effect of elevation on site occupancy estimates of *Iguana delicatissima* based on transect surveys conducted on St. Eustatius in 2017 (pre-hurricane), 2018 and 2019 (post-hurricane). Data obtained using R package *unmarked*.

**Figure S2.**
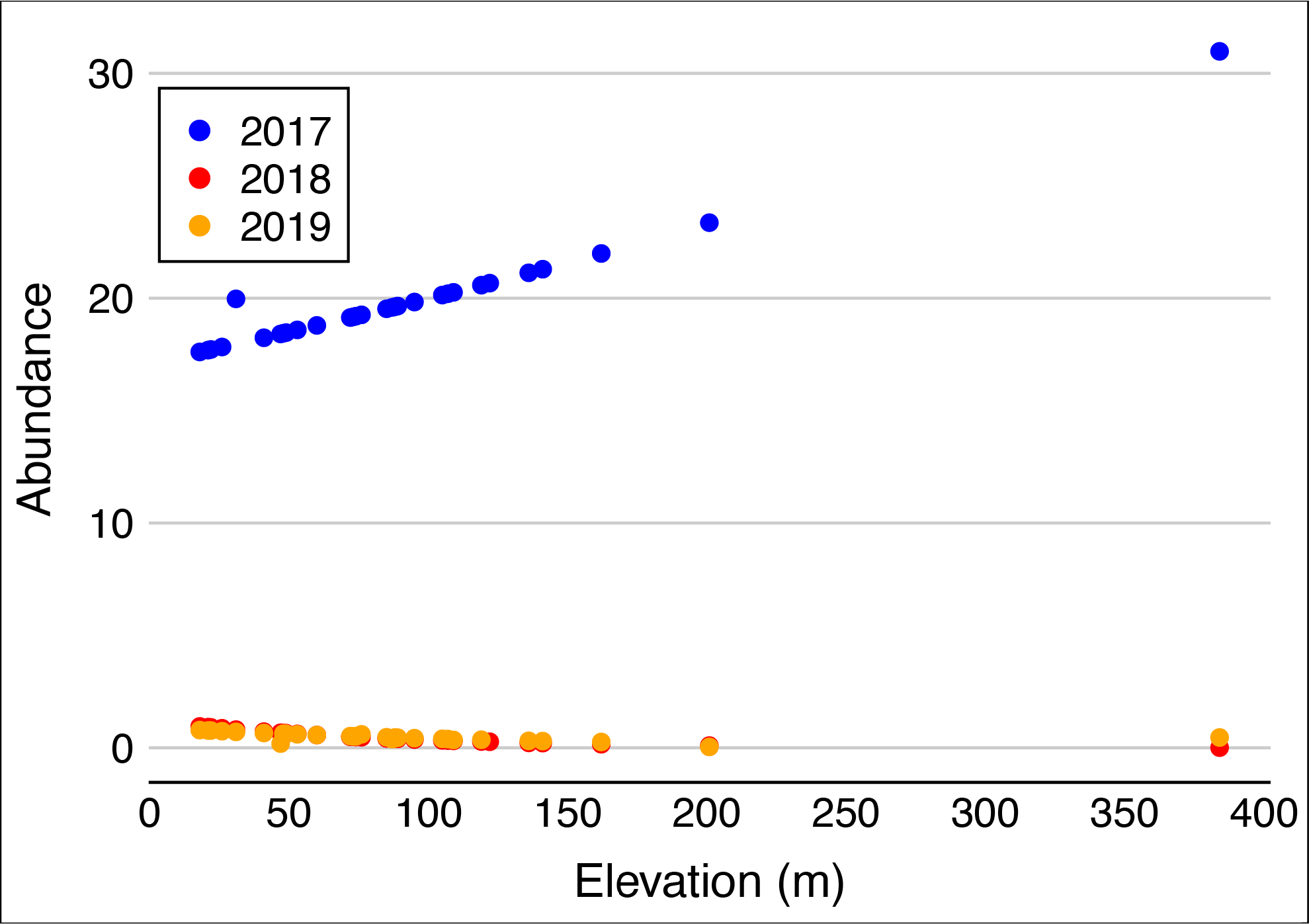
The effect of elevation on abundance estimates of *Iguana delicatissima* based on transect surveys conducted on St. Eustatius in 2017 (pre-hurricane), 2018 and 2019 (post-hurricane). Data obtained using R package *unmarked*.

**Figure S3.**
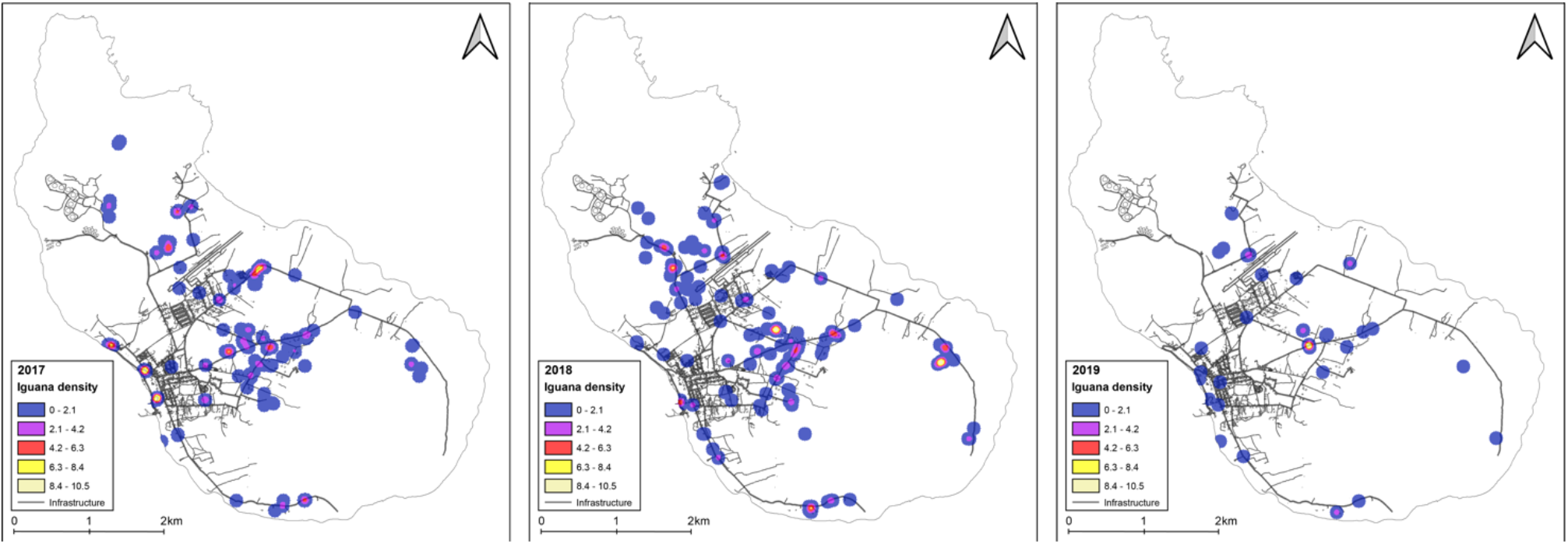
Iguana density heatmap for opportunistically observed bead-tagged iguanas on St. Eustatius, during 2017–2019.

**Table S1.**
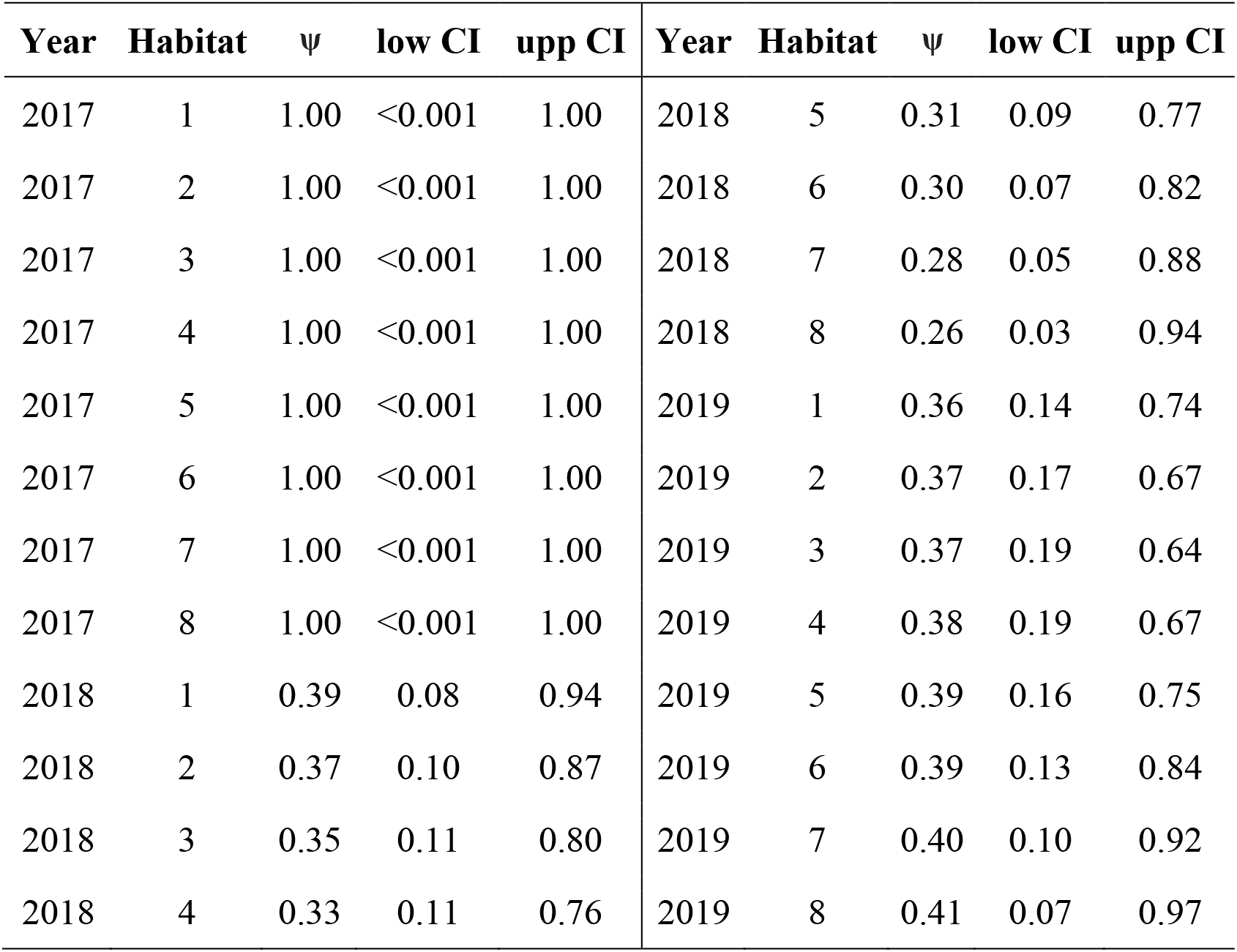
Overview of 2017–2019 site occupancy estimates per habitat of *Iguana delicatissima* on St. Eustatius. Ψ presents the probability of a site to be occupied, including lower and upper confidence intervals (CI). Habitat following De Freitas et al. (2012): 1 = *Pisonia* - *Antirhea* mountains; 2 = miscellaneous urban/agricultural or disturbed land; 3 = *Pisonia* - *Justicia* hills; 4 = *Capparis* - *Pisonia* mountains; 5 = *Rauvolfia* - *Antigonon* mountains; 6 = *Coccoloba* - *Bothriochloa* cliffs; 7 = *Pisonia* - *Eugenia* mountains; 8 = *Chionanthus* - *Nectandra* mountains.

| Year | Habitat | $\psi$ | low CI | upp CI | Year | Habitat | $\psi$ | low CI | upp CI |
| --- | --- | --- | --- | --- | --- | --- | --- | --- | --- |
| 2017 | 1 | 1.00 | <0.001 | 1.00 | 2018 | 5 | 0.31 | 0.09 | 0.77 |
| 2017 | 2 | 1.00 | <0.001 | 1.00 | 2018 | 6 | 0.30 | 0.07 | 0.82 |
| 2017 | 3 | 1.00 | <0.001 | 1.00 | 2018 | 7 | 0.28 | 0.05 | 0.88 |
| 2017 | 4 | 1.00 | <0.001 | 1.00 | 2018 | 8 | 0.26 | 0.03 | 0.94 |
| 2017 | 5 | 1.00 | <0.001 | 1.00 | 2019 | 1 | 0.36 | 0.14 | 0.74 |
| 2017 | 6 | 1.00 | <0.001 | 1.00 | 2019 | 2 | 0.37 | 0.17 | 0.67 |
| 2017 | 7 | 1.00 | <0.001 | 1.00 | 2019 | 3 | 0.37 | 0.19 | 0.64 |
| 2017 | 8 | 1.00 | <0.001 | 1.00 | 2019 | 4 | 0.38 | 0.19 | 0.67 |
| 2018 | 1 | 0.39 | 0.08 | 0.94 | 2019 | 5 | 0.39 | 0.16 | 0.75 |
| 2018 | 2 | 0.37 | 0.10 | 0.87 | 2019 | 6 | 0.39 | 0.13 | 0.84 |
| 2018 | 3 | 0.35 | 0.11 | 0.80 | 2019 | 7 | 0.40 | 0.10 | 0.92 |
| 2018 | 4 | 0.33 | 0.11 | 0.76 | 2019 | 8 | 0.41 | 0.07 | 0.97 |

Mean detection probability (*p*) per transect was 0.01 (inestimable CIs) using “unmarked”. Occupancy probability was 33.1% (11.4–73.8%) in 2018, and 37.5% (19.4–64.0%) in 2019. *p* was 0.40 (0.12–1.34) in 2018, and 0.47 (0.22–1.02) as estimated using “unmarked”.

## Notes

### Competing Interest Statement

The authors have declared no competing interest.

### Summary of Updates

Our previous version missed the supplementary results which have now been added

